# Distant water industrial fishing in developing countries: A case study of Madagascar

**DOI:** 10.1101/2021.05.13.444019

**Authors:** Easton R. White, Merrill Baker-Médard, Valeriia Vakhitova, Samantha Farquhar, Tendro Tondrasoa Ramaharitra

## Abstract

As industrial vessels continue to expand in both extractive capacity and spatial range, concerns have grown over foreign industrial fishing occurring within the marine territories of developing countries, both legally and illegally. Madagascar’s status as a “least developed country”, coupled with its high marine biodiversity, makes its waters particularly susceptible to fishing by distant water fishing nations (DWFNs). However, given constraints in management and research, it is unclear how foreign industrial fishing, both legal via foreign agreements and illegal, may impact local marine resources that many coastal communities depend on for food security, cultural meaning and livelihoods. We used satellite-derived fishing effort data from 2012-2020, via Global Fishing Watch, to analyze industrial fishing effort occurring within Madagascar’s Exclusive Economic Zone (EEZ). We documented 907,643 hours of industrial fishing within the Madagascar EEZ across 277 vessels from 17 different countries. We found that Taiwanease vessels (39.8%) using drifting longlines and Malagasy (17.2% shrimp trawlers were the most prevalent. Fishing effort was highly seasonal (68% of effort between October and February) and increased with higher global fish prices and the Indian Ocean Dipole, which is a measure of regional water temperature cycles. We also found a number of instances (17.6% of the fishing effort for 170,726 total hours) of foreign fishing vessels operating close to shore and within a number of marine protected areas. These results highlight the need for increased transparency surrounding foreign fishing agreements and unauthorized fishing within the waters of developing countries. Increases in industrial fishing effort and encroachment into near-shore areas has the potential to severely threaten current sustainable fisheries management initiatives by conservation organizations and coastal communities.

**Highlights:**

1. Distant water fishing nations dominated fishing efforts within Madagascar’s EEZ.
2. Longlining by foreign nations was the dominant fishing mode and increased from October-February.
3. Malagasy vessels focused on trawling for shrimp.
4. Fishing effort increased during positive Indian Ocean Dipoles and with higher fish prices.
5. Distant water fishing nations frequently fished close to shore and sometimes within MPAs.

## 1 Introduction

The range and capacity of industrial fishing has increased dramatically in the last century (Tickler et al., 2018). Industrial fishing now occurs in over 55% of our oceans (Belhabib et al., 2020; Kroodsma et al., 2018). This increase is driven by growing demand for marine resources as well as technological advances in industrial fishing (Galbraith et al., 2017). Although many fisheries are sustainably managed (Hilborn et al. 2020), declines of some fisheries have driven industrial fishing efforts far beyond a country’s own waters (Kroodsma et al., 2018; Pauly et al., 2014; Pauly & Zeller, 2016).

In many low-income countries, distant water fishing nations (DWFN) are a large and growing subset of a country’s industrial fishing profile (Nichols et al. 2015). For many DWFN, there is a high degree of illegal, unreported and unregulated (IUU) fishing that occurs, undermining efforts to achieve sustainable yield and manage local and regional fish stocks (Petrossian, 2015). These fleets, also known as “roving bandits,” account for an estimated 78% of industrial fishing effort occurring within the national waters of lower-income countries (McCauley et al., 2018, Berkes, 2006). When DWFN negotiate legal access to low-income countries EEZs, many licensing agreements have been shown to be predatory, undercutting the financial benefit and food security interests of the host country (Le Manach 2012, Gagern & van den Bergh, 2013).

Numerous studies report DWFN fishing in off-limit areas (Belhabib et al., 2020; Kimani et al., 2009), including marine protected areas (MPAs) (Pauly et al., 2014; T. D. White et al., 2020). Given that MPAs are considered the cornerstone of marine conservation (Edgar et al., 2014; Mellin et al., 2016; E.R. White et al., 2015), incursions in these areas by foreign industrial fishing fleets can undermine key management and conservation objectives by contributing to local fisheries collapse and causing cascading effects throughout the ecosystem (Cohen et al., 2019). Notable cases of illegal industrial fishing inside MPAs by DWFN include the Galapagos Marine Reserve (Carr et al., 2013), Cocos Island (E.R. White et al., 2015), and Papahānaumokuākea Marine National Monument of Hawai’i (T. D. White et al., 2020). However, most of this work has focused on large marine protected areas or countries with more management capacity. There is a lack of information concerning industrial fishing within MPAs in most countries in Sub-Saharan Africa (Belhabib et al., 2020).

Currently, the precise impact of foreign industrial fishing on local marine resources and resource users is not well understood. Obtaining accurate information concerning where, how, and when industrial fishing vessels operate is elusive to many governmental and non-governmental organizations (Cabral et al., 2018). Past work on the drivers of fluctuations in fishing effort over time (Stephenson et al., 2018), has demonstrated the importance of global oil prices and seafood prices as two important predictors of global fishing effort (Cheilari et al., 2013; Dahl & Oglend, 2014; Guillotreau et al., 2017; Kroodsma et al., 2018). In addition, biological productivity, large scale climate oscillations (e.g., El Niño, Indian Dipole Index), and extreme weather events have all been shown to be important predictors of commonly-fished pelagic species occurrence (Osgood et al., 2021) and fishing effort (Belhabib et al., 2018; Kroodsma et al., 2018; Kumar et al., 2014). However, this past work has typically focused on regional and global data sets, with implications for specific countries less clear (Kumar et al., 2014). This is largely due to the difficulty in obtaining accurate industrial fishing data, which is an especially pronounced challenge in a developing country context given funding and capacity constraints (Erceg, 2006; Kimani et al., 2009). Knowledge of the characteristics, location and frequency of fishing in a developing country where the majority of industrial activity is conducted by a foreign fleet is critical to both fisheries management as well as domestic food security and sovereignty (Le Manach et al., 2012).

In many ways, Madagascar is an ideal case study to analyze the complex social and ecological stakes of how and where industrial fishing occurs globally. Madagascar, classified as a “least developed country” by the United Nations, is also considered one of the most marine resource rich countries globally (Selig et al., 2014). It is estimated that the per capita gross domestic product (GDP) is 449.72 USD with the majority of the population (70 percent) living below the poverty line. This economic poverty, however, is coupled with immense marine natural resource wealth. The World Bank recently estimated that the value of the Gross Marine Product (GMP) in the Western Indian Ocean to be US$20.8 billion annually, making it the 4^th^ largest economy in the region in which over half was attributed to Madagascar (Obura, 2017). Madagascar, similar to many other developing countries with large EEZs, sells access to marine resources through permits to a large number of DWFN (Gagern & van den Bergh, 2013; McCauley et al., 2018, Le Manach et al., 2012). In 2018, the fishing industry constituted 7% of national GDP (World Bank, 2020).

Given the lack of transparency of many foreign fishing agreements coupled with inadequate monitoring and enforcement capacity within Madagascar, industrial fishing effort is difficult to determine in Madagascar. While Madagascar does have a Bureau of Fisheries Surveillance, its capacity to carry out its functions across Madagascar’s vast seascape is low. The office only employs 45 people and has only eight surveillance boats and ten motorboats (Antsa, 2020; Baker-Medard & Faber, 2020; Orange, 2021). A number of indices and scorecards have tried to demonstrate Madagascar’s vulnerability to IUU fishing. For example, the IUU Index provides a measure between 1 and 5 (1 being the best, and 5 the worst) of the degree to which states are exposed to and effectively combat IUU fishing. Madagascar received an IUU fishing coastal vulnerability score of 4.33 (Macfadyen et al., 2019).

Madagascar is also home to thousands of coastal communities who depend on marine resources for food and livelihood. An estimated 1.5 million people are dependent on fisheries and aquaculture in Madagascar with the majority from rural coastal areas (World Bank, 2020). This number is likely increasing given that more people are moving to the coast and pursuing fishing related livelihoods (Cripps & Gardner, 2016). Consequently, fisheries sustainability is viewed as a pressing problem in Madagascar as fish stocks are declining and coastal communities are some of the most vulnerable to food insecurity (Le Manach et al., 2012). Like many developing countries, Madagascar does not have large industrial fleets, instead the majority of fisheries production comes from small-scale fishers. Encroachment on nearshore fishing zones from industrial vessels has the potential to undermine this local provision of nutrient-rich food. One remedy the Malagasy government and international conservation organizations have implemented to help protect marine biodiversity and sustainably manage fisheries is the establishment of MPAs. Madagascar committed to drastically expand the marine area under protection both at the World Parks Congress in 2003 and through its commitments to Aichi Targets within the Convention on Biological Diversity in 2014 (SAPM 2009, CBD 2016).

The vast majority of MPAs established in Madagascar to date occur within three nautical miles of shore. Small-scale fishers (SSF) operate close to shore adjacent to these MPAs, while all foreign industrial vessels should legally operate outside territorial waters (12 nautical miles from the coast), or if unpermitted, outside the exclusive economic zone (EEZ, extending 200 nautical miles from the coast) (Randrianarisoa 2006, Bonzon 1986, Loi 85-013 of 11 December, 1985). In addition to the broad 12 nautical mile restriction on all foreign industrial vessels, there are additional restrictions on domestic tuna fishing within 8 nautical miles and all domestic shrimp trawling within 2 nautical miles of shore to protect small-scale fishers (Randrianarisoa 2006). With increased adoption of automatic identification systems (AIS) using satellite tracking, there is growing evidence that illegal industrial fishing occurs frequently nearshore. For example, Belhabib et al. (2020) found that industrial fleets spend 3%–6% of their time fishing within nearshore areas reserved for SSF across Africa. Similarly, Malagasy SFF in multiple regions of the island have reported foreign industrial fishing vessels encroaching in their coastal fishing areas (Baker-Médard, pers. obs.).

In order to address the challenges of data limitations associated with industrial fishing, we used remotely-sensed estimates of fishing effort. Specifically, we used data from Global Fishing Watch (Global Fishing Watch, 2021; Kroodsma et al., 2018) to assess the spatio-temporal dynamics of industrial fishing within Madagascar’s EEZ from 2012-2020. These data include daily locations of inferred fishing activity (Global Fishing Watch, 2021). Specifically, we address four questions: 1) Who is fishing in Malagasy waters and where are they fishing?; 2) How has fishing effort changed over time?; 3) What are the best predictors (e.g., economic versus environmental) of fishing effort?; and 4) How much fishing has occurred nearshore and within protected areas? We hypothesized that fishing effort would increase over time by distant foreign nations, that fishing would be correlated with sea surface temperature and seasonality, and lower cyclone activity. In addition, we hypothesized that fishing effort would increase with fish prices and decrease with oil prices. Lastly, we hypothesized that most fishing would be away from outside of nearshore or protected areas.

## 2. Methods

### 2.1 Fishing effort data

We used data from Global Fishing Watch (GFW) from January 2012 to December 2020. Global Fishing Watch uses vessel movement data from Automatic Identification System (AIS) transmissions and a set of machine learning approaches to distinguish general vessel movement from species types of fishing (Kroodsma et al., 2018). The resulting records include information on the daily fishing effort represented by hours spent in a perimeter gridded by 0.1° of longitude and latitude, flag state, gear type, and vessel information (e.g., size, registration) (Kroodsma et al., 2018). Details on this dataset are best found in Kroodsma et al., 2018 and on the Global Fishing Watch website. In order to focus on the Malagasy EEZ, we selected the GFW records present only within the Malagasy EEZ. This resulted in an abridged database containing records only for 277 vessels with gross tonnage varying from 19 to 3674 tons in volume. This consists of vessels larger than 15 meters.

### 2.2 Environmental and economic covariates

We examined several environmental and economic covariates as potential explanatory variables for fishing effort (Table 1). We used an index of the Indian Ocean Dipole Mode (NOAA, 2020; Saji et al., 1999) as a measure of anomalous difference in sea surface temperature (SST) between the western (50°E to 70°E and 10°S to 10°N) and eastern (90°E to 110°E and 10°S to 0°S) Indian Ocean (*Bom.Gov.Au*, 2020). A positive dipole score indicates a period of warmer waters, accompanying wind and increased precipitation in the Western Indian Ocean, whereas negative score signifies anomalously low SST around Madagascar (Cai et al., 2014) that is known to cause drought in eastern Africa (Lim & Hendon, 2017).

**Table 1.** Explanatory variables used as fixed effects in models of industrial fishing effort from 2012-2020 around Madagascar.

| Variable | Range | Description | Source |
| --- | --- | --- | --- |
| Year | 2012-2020 |  |  |
| Julian day | 1-365 | Day of the year |  |
| Dipole index | (-1.37) –<br>(+2.15) | Indian Ocean Dipole (IOD) is an index that shows the changes in the difference between the sea surface temperature of the tropical western and eastern Indian Ocean. Positive numbers correlate with weakened westerly winds and warmer than average waters shifting towards Africa (50°E to 70°E) | 2021 Australian Bureau of Meteorology (*Bom.Gov.Au*, 2020) |
| Global oil price | (-54.2942)<br>–<br>(248.1612) | USD/bbl | (“IMF Primary Commodity Prices,” 2020) |
| Global fish prices | 4.35 - 8.06 | USD per kilogram | (“IMF Primary Commodity Prices,” 2020) |
| Cyclone events | Presence or absence on any given day | Presence of any cyclone within the Madagascar EEZ with a cyclone defined as any form of sustained wind speed above 15 knots (Tropical Cyclone Climatology 2015) | NOAA IBTrACS Version 4 |

We used data from NOAA IBTrACS Version 4 to obtain a binary variable for the presence or absence of cyclone activity within the South Indian Ocean (Knapp et al., 2018) from 2012-2016 and further limited data to the Madagascar’s EEZ. If a severe storm event received a name, we considered it a cyclone. A cyclone in this data is defined as any form of sustained wind speed above 15 knots (*Tropical Cyclone Climatology*, 2015).

We also included global oil and fish prices as economic indicators. World crude oil prices and fish prices were taken from International Monetary Fund (IMF) Data. We considered monthly values of oil prices in USD units. Average petroleum spot price (ASPS) of crude oil calculated by IMF was presented in $/bbl (*IMF Primary Commodity Prices*, 2020) and derived from the simple average of three spot prices; Dated Brent, West Texas Intermediate, and the Dubai Fateh (*IMF Primary Commodity Prices*, 2020). As a measure of global fish prices, we used the monthly export price of farm-bred Norwegian salmon in USD per kilogram for the 2012-2019 period (*IMF Primary Commodity Prices*, 2020). Although it is not an ideal measure for species caught around Madagascar, it is highly correlated with other seafood products given the globalized nature of much of the industry (*IMF Primary Commodity Prices*, 2020).

### 2.3 Modeling approach and analysis

We used generalized linear models with a Gaussian error distribution (i.e., linear regression) to assess the predictors of fishing effort. Because the number of vessels included in the database changed during the course, we decided to run models only for the vessels that were present in all nine years of the study (n=51). To test the sensitivity of our results, we examine other models in the supplementary material that focus on either only recent (and higher quality) data or fishing effort that has been standardized by the number of observed vessels. We first examined the total fishing effort (in terms of hours) per day for the entire Madagascar EEZ for 2012-2019. We did not include 2020 in these models given missing data for the covariates. In addition to environmental and economic explanatory variables (Table 1), we included sine and cosine functions of Julian date as explanatory variables to account for seasonality (Baum & Blanchard, 2010; E.R. White et al., 2015). We examined all potential combinations of explanatory variables (n = 96) using the MuMIn package (Bartoń, 2013). We did not include combinations of explanatory variables that were highly (>0.7) correlated to account for issues of multicollinearity (Vatcheva et al., 2016). We then selected the best fitting model or if several models fell within 2 Akaike information criterion (AIC) of the top model, we used a model averaging approach (Zuur et al., 2009) to generate parameter estimates. We verified model assumptions by examining plots of the Pearson residuals (Fig. S7). All models were implemented in R (R Core Team, 2020).

### 2.4 MPA and LMMA database

In order to create a master database and shapefile for all MPA’s and Locally Managed Marine Areas (LMMA’s) on Madagascar we consulted multiple resources including both previously established databases, personal observations and literature. We used Systèm des Aires Protégée de Madagascar (*Rebioma*, 2017) information and shapefiles as the foundation of the map. We further added missing protected areas from Madagascar National Parks (*Madagascar National Parks*, 2020), UNESCO World Heritage Sites (UNESCO, 2020), WCS Madagascar (*WCS Madagascar*, 2020), Ramsar Wetlands of International Importance (*Madagascar Ramsar*, 2020) and The World Database on Protected Areas (*UNEP-WCMC*, 2020). Every MPA and LMMA listed in Table S4 was verified by at least two references in journals or NGO publications. In the case of Anoromby/Andreba LMMA presented by Butler, 2015 no available references were found, as a result we referred to personal communication with Dr. Baker-Medard for verifying the veracity of the information.

The MPA and LMMA polygons missing from the resulting shapefile but mentioned in literature were drawn manually to resemble the documented area. For instance, Antongil Bay Shark Sanctuary was depicted based on the article published in Mongabay News (Butler, 2015). Sainte Luce No Take Zone (NTZ), Elodrato NTZ, Itapera NTZ and Sainte Luce LMMA were created based on previous work (Long et al., 2019).

## 3 Results

### 3.1 Temporal dynamics

Between 2012-2020, there were 907,643 documented hours of fishing within the Madagascar EEZ. This does not include all industrial fishing efforts as vessels were added to the database during the course of the study and some vessels were likely not included. In total, there were 277 vessels recorded fishing from a total of 17 different countries (Fig. 1). Taiwan accounted for 39.8 percent of all fishing effort, with other distant water fishing nations (France, Japan, China, Korea, Malaysia, and Spain), constituting the bulk of other longlining activity (Fig. 1). Madagascar accounted for 17.2 percent of all fishing effort, led by shrimp trawlers, which were first recorded in 2018, although they were present before then (Farquhar, Baker-Médard, pers. obs.). Greek vessels were the only other major source of trawling within the EEZ (Fig. 1).

**Figure 1.**
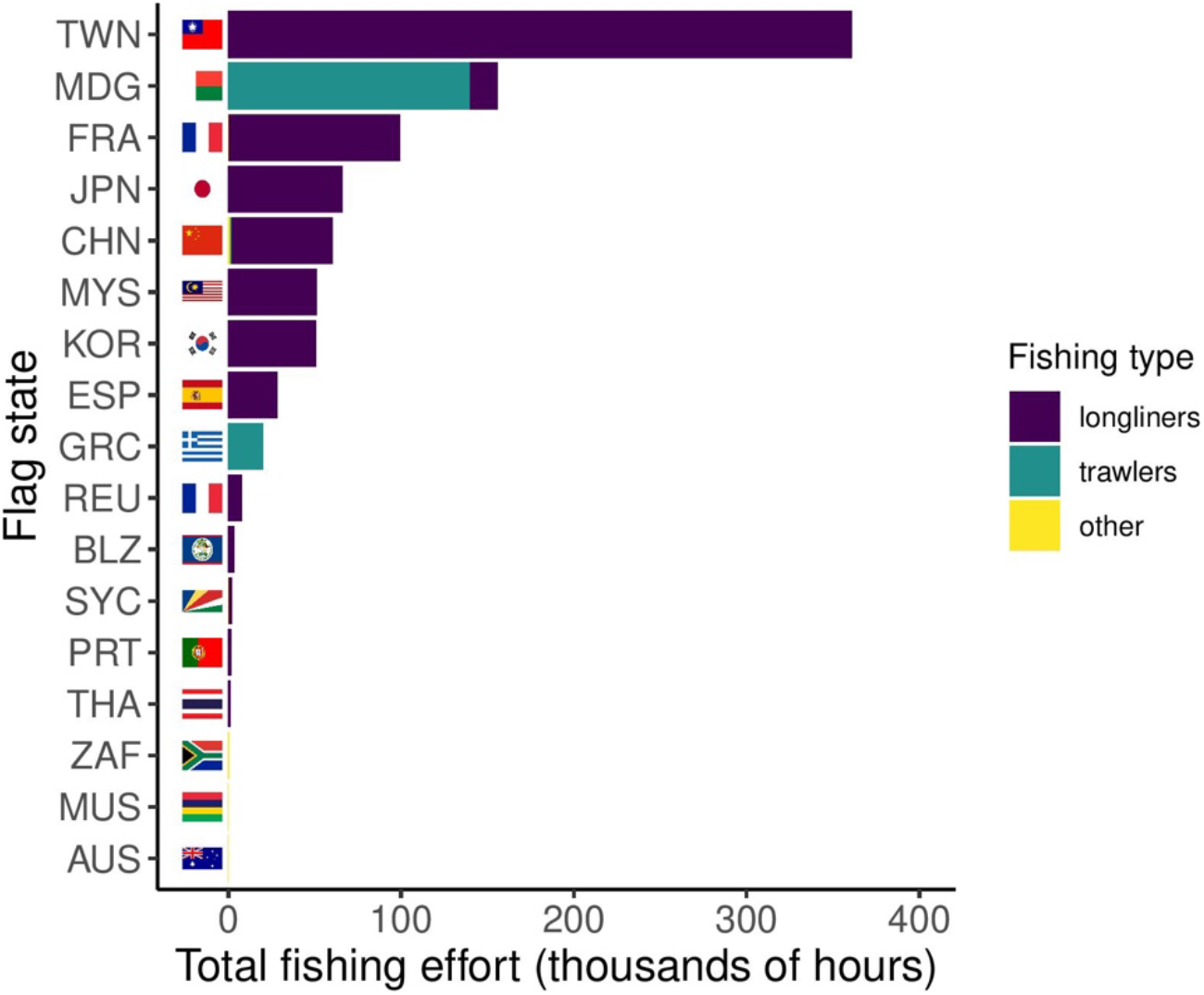
Total fishing effort (in hours) for the entire Madagascar EEZ from 2012-2020 by flag state and gear type. The other gear type designation refers to not-descriptive fishing, pole and line, set gillnets, squid jiggers, and tuna purse seiners.

The vast majority of fishing activity was drifting longlines (82.1%) and trawlers (17.7%) with vessels mostly in the 25-30 meter and 50-150 tonnage range (Figs. 1, S1). Fishing effort was highly seasonal and peaks between November-January each year (Fig. 2). When standardized by cumulative number of vessels recorded, fishing effort was similar between years (Fig. 2). Thus, given the nature of the data, it is not possible to determine if the total fishing effort is truly changing over time or if it is actually the number of vessels included in the data (Fig. 2). For example, in years 2017-2020, the data showed that there were many Madagascar flagged vessels operating in nearshore areas, including protected areas, on the west coast. These are almost exclusively part of the national shrimp trawling fleet who have been active since the 1960s (Le Manch *et a. 2012).* However, they only likely began appearing in the data as the fleet has been modernized with AIS.

**Figure 2.**
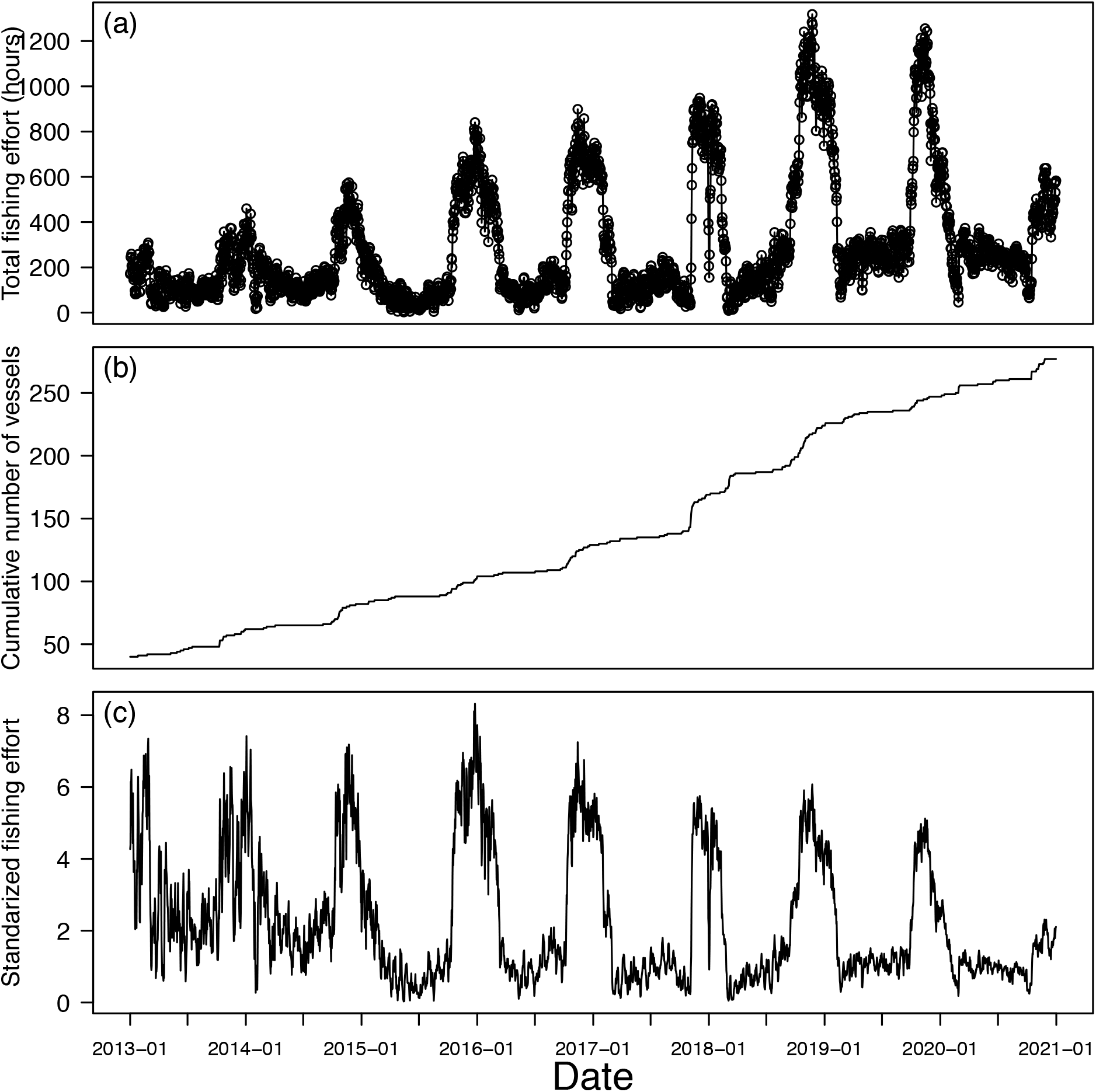
(a) Total fishing effort within the Madagascar EEZ per day. (b) Cumulative number of vessels observed within the Madagascar EEZ per day. (c) Standardized fishing effort which is the total fishing effort (in hours) divided by the cumulative number of vessels observed. Effort from 2012 not shown due to low number of vessels recorded.

### 3.2 Temporal covariates

In addition to being highly seasonal (significance of sine and cosine terms in Table 2), fishing effort was also strongly correlated with a number of economic and environmental covariates (Table 2). We found that fishing effort was higher with a positive dipole index, indicating more fishing during periods of warmer waters, increased wind speeds and increased precipitation (Table 2, Figs. S6). Fishing effort was not strongly correlated with the presence of cyclone events (Table 2, Fig. S5). In addition, fishing effort increased with higher fish prices (Table 2, Fig. S4). Conversely, fishing effort decreased with higher global oil prices (Table 2, Fig. S3)

**Table 2.** Parameter estimates (top value) and standard errors (bottom value in parentheses) for the average of the best fitting models (AIC < 2) using a GLM framework with a Gaussian error structure. Fishing effort (from 2012-2019) is the total daily fishing effort within the Madagascar EEZ for only the vessels seen during all nine years of the data set (n = 51). Sine and cosine terms denote seasonal variables as a function of the Julian day of the year.

|  | **Fishing effort** |
| --- | --- |
| (Intercept) | 27115.7***<br>(1475.3) |
| sine | -22.9***<br>(1.4) |
| cosine | 45.1***<br>(1.3) |
| year | -13.5***<br>(0.7) |
| Dipole index | 28.6***<br>(4.1) |
| Fish price index<br>(USD) | 20.4***<br>(1.1) |
| Oil price (USD) | -0.36***<br>(0.1) |
| Cycle event | -2.3<br>(4.6) |

\*\*\* $p < 0.001$ ; \*\* $p < 0.01$ ; \* $p < 0.05$

### 3.3 Spatial dynamics

The exact distribution of fishing effort changed over time, fishing was generally concentrated on the east coast, south of the island, and on the western coast between Morondava and Mahajanga (Fig. 3). Most of the 2012-2020 fishing effort (82.4%) was 12 nautical miles (22.2 km) or more from shore, however, 17.6% of the fishing effort was closer to shore (Figs. 3,4). This accounts for approximately 170,726 total hours of fishing during the course of the study or 52.7 hours of fishing effort per day. In addition to fishing effort nearshore, we also documented fishing vessels fishing within marine protected areas (Fig. 4). As a pair of case studies, we chose two marine protected areas, Barren Isles Archipelago and Ambodivahibe. Barren Isles is the largest marine and coastal protected area and was given temporary status in 2014, and official status in 2017. Our analysis found that between 2013-2020, multiple trawling vessels flagged to Greece operated in the Barren Isles MPA. The Ambodivahibe protected area was first established in 2009 and includes an area of 465.62 km^2^. Drifting longline and purse seiners vessels from Seychelles, France and Reunion were observed in this area between 2013-2020. It should be noted that other protected areas had similar incursions such as the Baie de Baly, Mahavavy Kinkony, and Ankivonjy.

**Figure 3.**
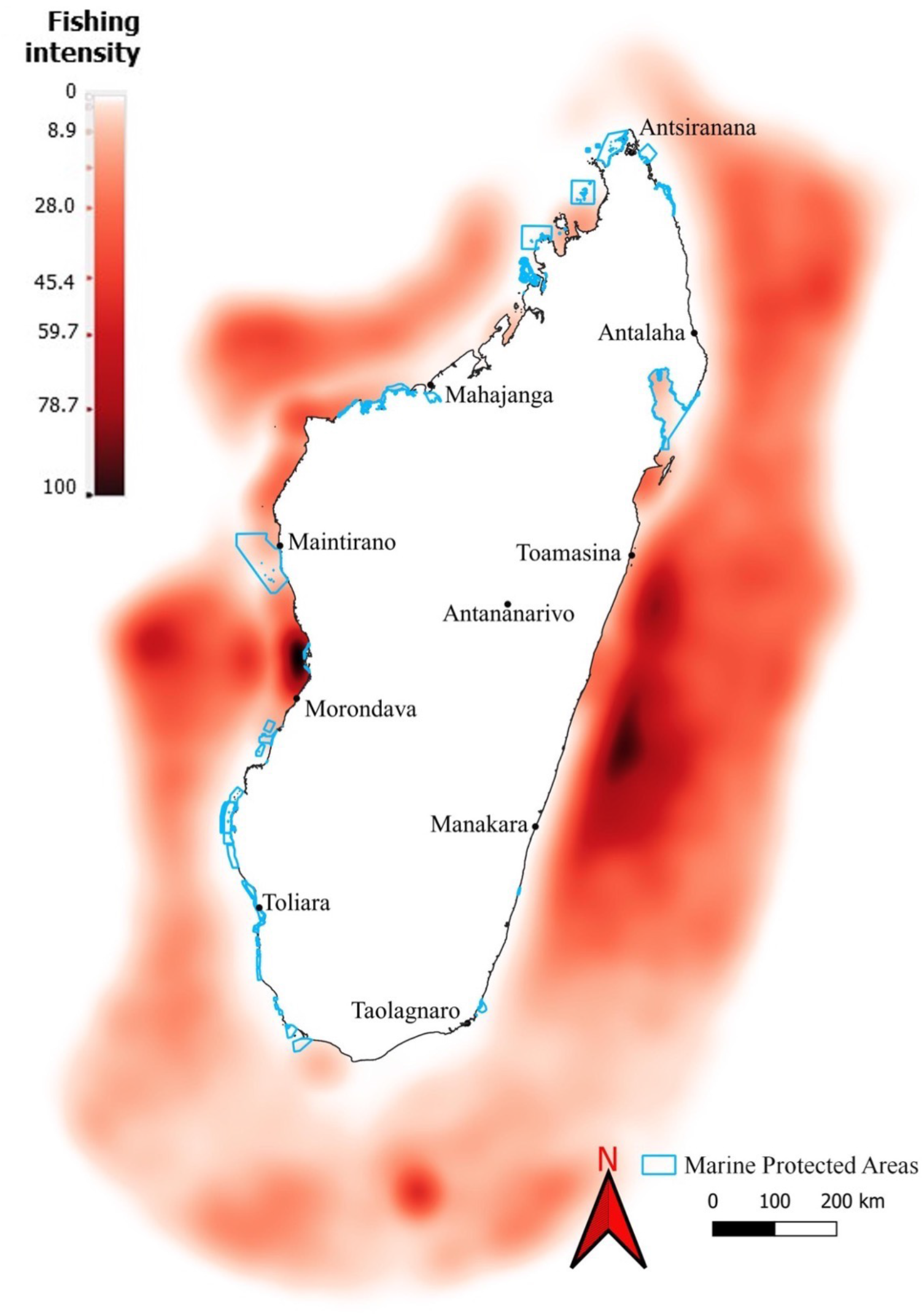
Total fishing effort standardized between 0 and 100 (a percentage of max effort) across the entire Madagascar EEZ from 2012-2020 with darker areas indicated more effort and lines denoting marine protected areas.

**Figure 4.**
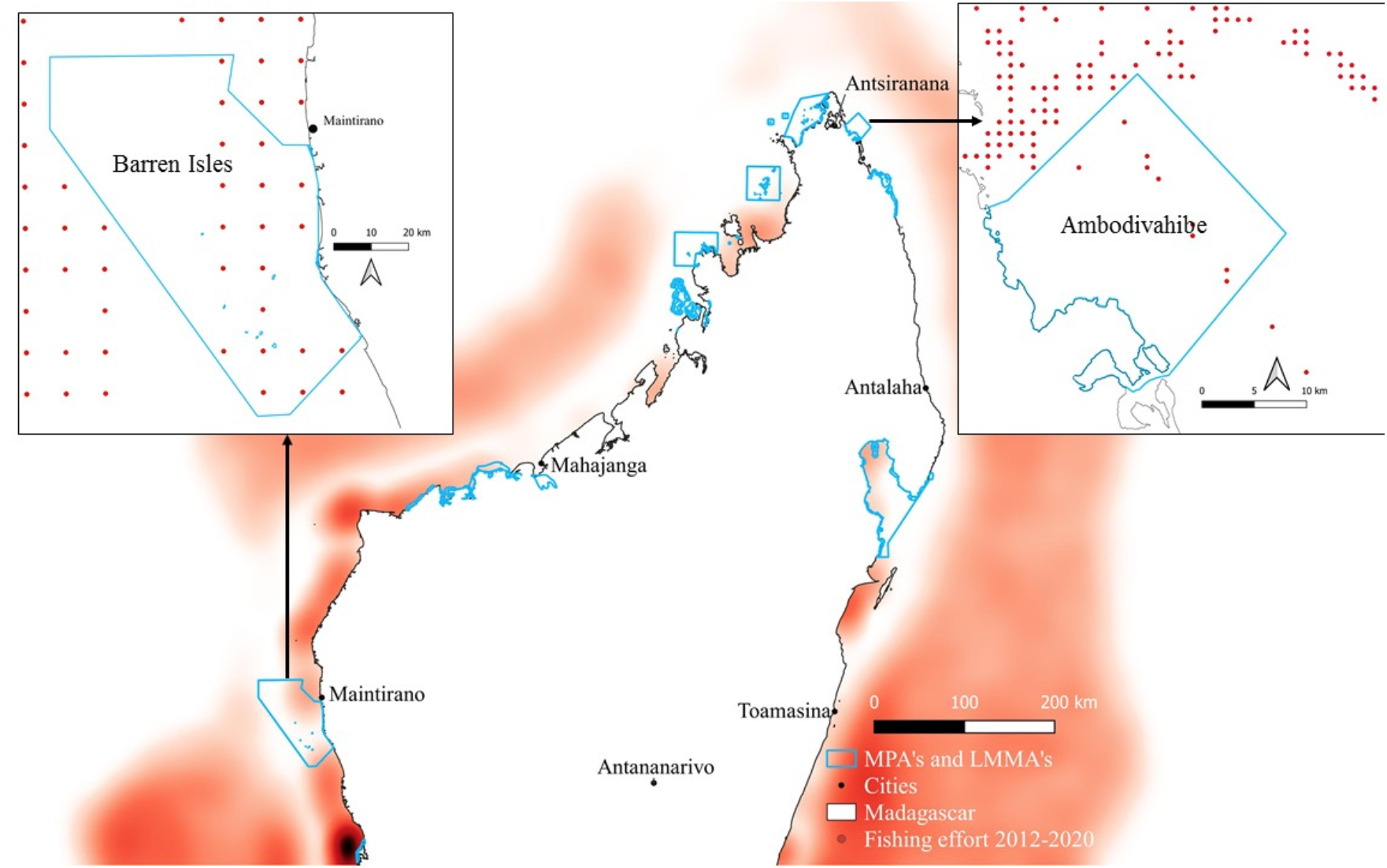
(larger map) Total fishing effort from 2012-2020 with darker areas indicated more effort and lines denoting marine protected areas. (smaller maps) Fishing vessel movement in two marine protected areas: Barren Islands and Ambodivahibe. The red dots indicate vessel locations.

## 4 Discussion

In developing countries, industrial fishing activity, especially by distant-water fishing fleets, is notoriously difficult to monitor and regulate. This is largely due to lack of financial, technical, and human capacity to fulfil such duties (Erceg, 2006). As a case study into these issues, we evaluated the industrial fishing activity within the Madagascar EEZ using inferred fishing effort data from Global Fishing Watch. From 2012-2020, we found 907,643 hours of fishing effort across 277 vessels throughout the Madagascar EEZ. In line with past work, we found that the most popular (82.1%) gear type of industrial fishing is drifting longlines, a method of fishing either pelagic or demersal species by comprising main lines to which additional lines are attached, each ending with baited or artificial lures (Kimani et al., 2009). Target species of longliners are primarily tuna, swordfish and sharks (FAO, 2020). The Indian Ocean represents 20-24 percent of the world tuna supply, amounting to $2.3 billion annually (Moustahfid et al., 2018). Trawling activity was the second most common (17.7%) form of fishing in the area (Fig. 1). Malagasy shrimp trawlers constituted most of this activity which confirms past work (Razafindrainibe, 2010; Rojat et al., 2004; Cooke, 1997).

DWF fleets dominated the industrial fishing activity within the Madagascar EEZ (Figs. 1, S1). Specifically, Taiwan accounted for a combined 39.8% of all fishing effort with France, Japan, China, Korea, Malaysia, and Spain contributing to additional longlining fishing effort (Fig. 1) This is in line with previous work documenting fishing effort in the region (Andriamahefazafy, 2019; Cooke, 1997). However, trawling vessels from Greece and Madagascar were found to be fishing the most in the nearshore area. Foreign vessels fishing with licenses or permits authorized by the Malagasy government are not illegal. While the Malagasy government signs fishing agreements with foreign countries, these licensing deals lack transparency and have caused conflict between Malagasy fishers and Ministry representatives in the past. Equity issues loom large with regards to predatory fishing agreements between wealthy DWFN and economically poor yet resource rich countries like Madagascar (Gagern & van den Bergh, 2013; Nichols et al., 2015). For example, Le Manach *et al.* 2012 analyzed fishing agreements between the European Union (EU) and Madagascar since 1986 and found that while EU quotas increased by 30% over time, Madagascar’s financial benefit from these agreements decreased by 90%. Similarly, news broke in 2018 regarding a 10-year, $2.7 billion fishing deal between the Agency for Economic Development and Business Promotion (AMDP) and a group of Chinese companies, allowing 330 Chinese fishing vessels to operate in Malagasy waters. This deal was met with immense domestic and international public backlash, forcing the deal to be scrapped and ultimately, the Minister of Agriculture, Livestock and Fisheries claimed not to know about it (Carver, 2018). However, an agreement was signed by the Ministry of Agriculture, Livestock and Fisheries of Madagascar (MAEPM) one year later in 2019 that gives access to 28 Chinese vessels to both nearshore and offshore areas (Anon., 2020). While China was not the dominant DWF nation in our dataset, their interest in countries like Madagascar and other coastal African nations continues to increase as China seeks new bilateral partnerships to secure access to marine resources (Mallory, 2013).

Due to data limitations, we were not able to determine if there was an increase or decrease in fishing effort over time (Fig. 2). However, the number of fishing vessels included in the database did steadily increase over time (Fig. 2b). This could be from a real increase in the number of vessels fishing in the area or simply an inclusion of more vessels into the AIS database. Regardless of how we standardized the fishing effort data, we found that fishing effort was highly seasonal (Fig. 2, Table 2). Although fishing effort was noted year-round, approximately 68% of fishing activity occurred between the months of October and February. This is in line with past work in the Western Indian Ocean (Guiet et al., 2019), which has identified pelagic species movement and human society activity (e.g., the annual Chinese fishing moratorium, societal holidays) to be the main drivers of this seasonality. The timing of this activity is similar to small-scale fishing effort in other areas of the Western Indian Ocean where January typically marks the peak in production (Kylao et al. 2013).

In addition to seasonality, we had a number of other hypotheses on how fishing effort might be correlated with environmental and economic factors. The Indian Ocean Dipole is a measure of temperature anomalies (Cai et al., 2014; Saji et al., 1999). We found that fishing effort was greater during positive dipole events (Table 2, Fig. S6). This implies that fishing effort increased when water temperatures were warmer and there was increased precipitation in Madagascar. Contrary to our initial predictions, fishing effort was not correlated with the presence or absence of cyclone activity (Table 2, Fig. S5). This may be due to the ability of large, industrial vessels to avoid these types of events (Belhabib et al., 2018). We hypothesized that fishing effort might be correlated with global oil and fish prices. In line with our predictions, fishing effort did decrease with increases in global oil prices (Table 2, Fig. S3). Previous work on the role of fuel costs on fishing effort has been mixed. Some analyses of European fleets suggest price sensitivity to changes in fuel prices (Cheilari et al., 2013). However, other work suggests that more globalized fleets may be less price sensitive given large fuel subsidies (Kroodsma et al., 2018). Lastly, we found that fishing effort did increase with increases in global fish prices (Table 2, Fig. S4). Generally, global fish prices are very volatile over time and differ between species and product forms (Dahl & Oglend, 2014). Given the amount of fishing effort by long-lining vessels in the Madagascar EEZ, it is reasonable to infer that most effort is targeting tuna and other large pelagics (Andriamahefazafy, 2019; Michael et al., 2017). Our results also align with previous work on tuna markets. Manufacturer demand for tuna is considered to be relatively price elastic (Guillotreau et al., 2017), including in the Indian Ocean (Garcia del Hoyo et al., 2010). This is particularly true for yellowfin tuna (*Thunnus albacares*) which dominate the Indian Ocean catch and are more valuable than many other species (Guillotreau et al., 2017).

Although the distribution of fishing effort changed from year to year, fishing was generally concentrated on the east coast, south of the island, and along the western coast between Morondava and Mahajanga (Fig. 3). This could be explained by various oceanographic factors unique to Madagascar. Off the coast of southern Madagascar is the Madagascar Plateau. This plateau interacts with the southern flowing East Madagascar Current which results in a number of eddies, high turbulence, upwelling and consequently, highly productive areas (DiMarco et al., 2000; Lutjeharms & Machu, 2000; Machu et al., 2002; Ramanantsoa et al., 2018). On the west coast, research has shown that the nearshore area has some of the highest primary productivity levels in all of Madagascar (Pripp et al., 2014). This could be driven by the nutrients brought by a number of rivers as well as circulation from the Mozambique channel eddies (José et al., 2014; Pripp et al., 2014). On the east coast, the majority of fishing effort was concentrated close to the continental shelf break. Continental shelves can be another source of upwelling as colder waters are driven up to the surface thereby increasing productivity.

While fishing by authorized foreign vessels in Madagascar is legal generally, we also documented fishing effort closer nearshore and within protected areas (Figs. 3,4). This confirms anecdotal reports from coastal community members in Madagascar who claim industrial fishing vessels “steal” their fish. (pers. comm. Baker-Médard). When foreign vessels fish nearshore there is potential for conflict with SSF (Belhabib et al., 2014; DuBois & Zografos, 2012). This includes threats to marine safety, local food security, and livelihoods. Coastal Malagasy communities especially on the western coast of Madagacar have expressed frustration that industrial vessels ruin their fishing gear as they often run over and cut their nets (pers. comm. Baker-Médard and Farquhar). A better understanding of the legal and social dynamics of foreign industrial encroachment in nearshore waters is needed. For the SFF communities in the region who have been actively working to secure rights and implement sustainable management policies for marine resources, such encroachment undermines their social and environmental gains.

There are several policy implications that emerge from our findings. The first is that AIS can be a useful tool to help monitor both permitted and illegal fishing. Utilizing remote sensing tools to monitor fishing activity is an important way to help counter the lack of financial resources and infrastructure available to resource-rich low-income nations with large EEZs such as Madagascar. AIS, including data from Global Fishing Watch have been used to successfully fine vessels engaged in illegal fishing in several African nations (Dunn et al. 2018). These efforts are tied to several initiatives such as FISH-i Africa, FishCRIME, and West Africa Task Force that focus on stopping illegal fishing across Africa (https://stopillegalfishing.com/). Madagascar is part of FISH-i Africa Task Force, a partnership between eight countries of the Western Indian Ocean (Comoros, Kenya, Madagascar, Mauritius, Mozambique, Seychelles, Somalia and Tanzania) that enables governmental authorities to identify and act against large-scale illegal, unregulated and unreported fishing. FISH-i Africa, established in 2012, relies on satellite tracking and data such as AIS to stop illegal catch getting to market, fine vessels engaged in illegal activity, and prevent illegal vessels from operating in partner country EEZs. Thus far, FISH-i Africa has successfully caught and fined vessels engaged in illegal activity resulting in numerous arrests and multi-million dollar payments to partner countries (FISH-i Africa, 2017; Dunn et al., 2018). In its first two years alone FISH-i Africa’s action against several notorious illegal fishing operators that circulate widely in the Western Indian Ocean resulted in 3 million USD collected in fines (FISH-i Africa, 2016).

The second policy implication that emerges from our findings is that AIS is a useful tool to help monitor small MPAs. While researchers have lauded AIS monitoring in large remote MPAs (McCauley et al., 2016; White et al., 2020), our findings indicate that even small nearshore MPA can benefit from AIS monitoring. Utilizing AIS data to monitor industrial fishing within nearshore MPAs will allow countries to improve marine biodiversity conservation efforts, reduce the impact of industrial fishing on small-scale fishers, and adjust permitting agreements with nations that disregard domestic MPA boundaries.

There are several limitations with our approach. Although AIS-derived fishing effort data has been used successfully in many contexts (Belhabib et al., 2020; Kroodsma et al. 2018), it still has several limitations that are particularly relevant in the western Indian Ocean and Madagascar. AIS information is still biased towards open waters, vessels from wealthier nations, and industrial fishing vessels (Kroodsma et al., 2018). In addition, AIS transponders can be turned off, reducing our ability to track nearshore and illegal activity by vessels in Madagascar. Although the algorithms used to classify fishing type have an overall accuracy over 90% (Kroodsma et al., 2018), there are misclassifications which can vary spatially. We were able to document that industrial fishing is happening with Malagasy MPAs, LMMAs, and nearshore. However, more detailed work is needed to understand the specific relationship between industrial fishing and Malagasy, small-scale fisheries. Although it is outside the scope of this work, future work should evaluate how fishing effort has changed more recently. This is especially relevant given changes in the Malagasy government, new fishing agreements, and the COVID-19 pandemic (Bennett et al., 2020; Nichols et al., 2015; White et al., 2021).

## 5 Conclusion

Ultimately, we provide a snapshot of the spatiotemporal aspects of foreign industrial fishing occurring in Madagascar. We documented that distant-water fishing fleets dominate Madagascar’s overall industrial fishing and that they often operate nearshore or in protected areas. We also found that fishing effort increased during positive Indian Ocean dipole periods and with higher fish prices. Additional work is needed to understand how these results may have changed more recently and how industrial, foreign fisheries interact with local small-scale fisheries. This is especially relevant in the context of how wealth and other factors dicate who is involved in marine management in Madagascar (Baker-Médard et al., 2021). In particular, a more detailed examination on the effectiveness of marine protected areas in the context of foreign industrial fishing is needed. As Madagascar, and other similar countries, continue to invest in its fisheries, both nationally and through foreign fishing agreements, it is important to understand how such investment might be impacting coastal communities. Deepening our analysis of both the presence and predictors of distant-water fishing effort in Madagascar and other developing nations will ultimately lead to more informed, sustainable and just marine resource management decisions.

## 6 Acknowledgements

The authors thank Jiaqi Li, David Kroodsma, Tyler Clavelle, and the Global Fishing Watch team for their helpful comments and feedback with data analysis.

## 8 Data and code availability

All data used in this study is publicly available (see Table 1 for references). The code for figures and analyses is available at https://github.com/eastonwhite/GFW-Madagascar

