## Supplementary Material for "Distant water industrial fishing in developing countries: A case study of Madagascar"

#### Vessel characteristics

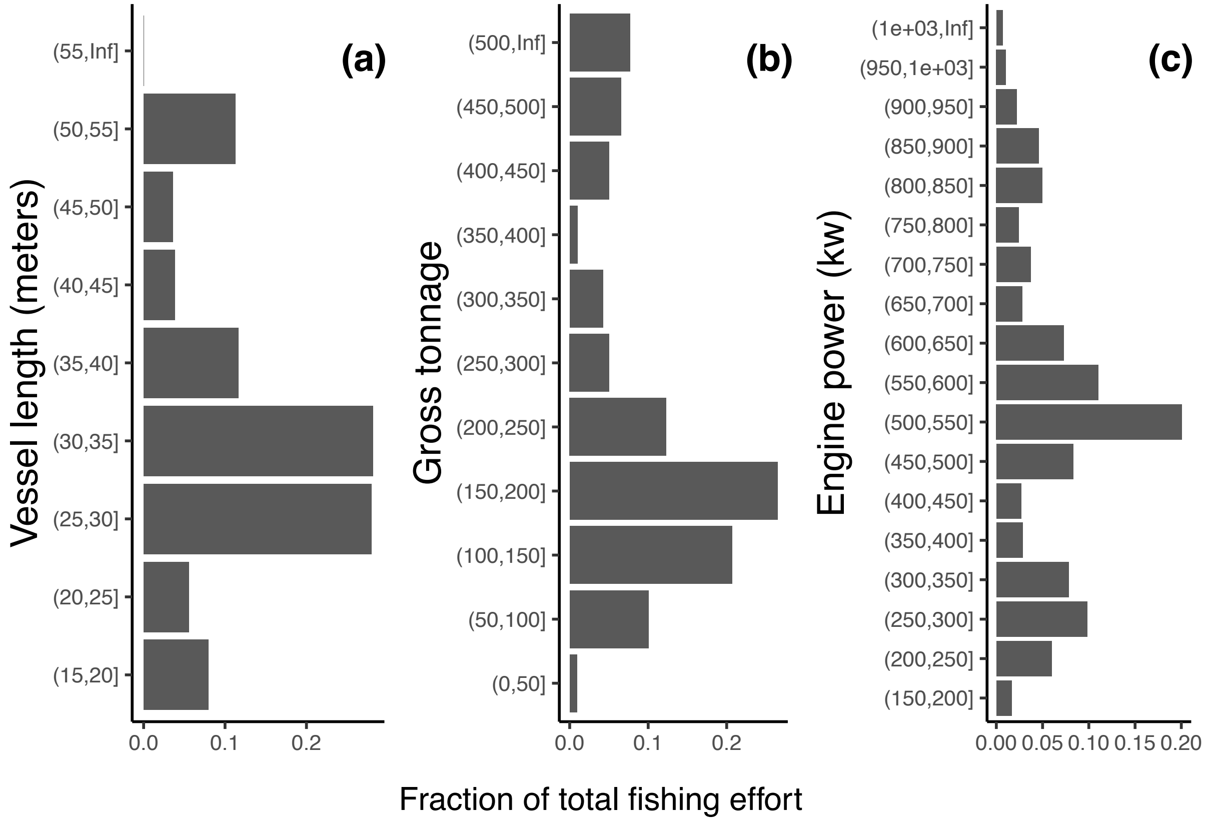


Figure S1: Total fishing effort (hours) versus various vessel characteristics: (a) vessel length (meters), (b) vessel tonnage, and (c) engine power

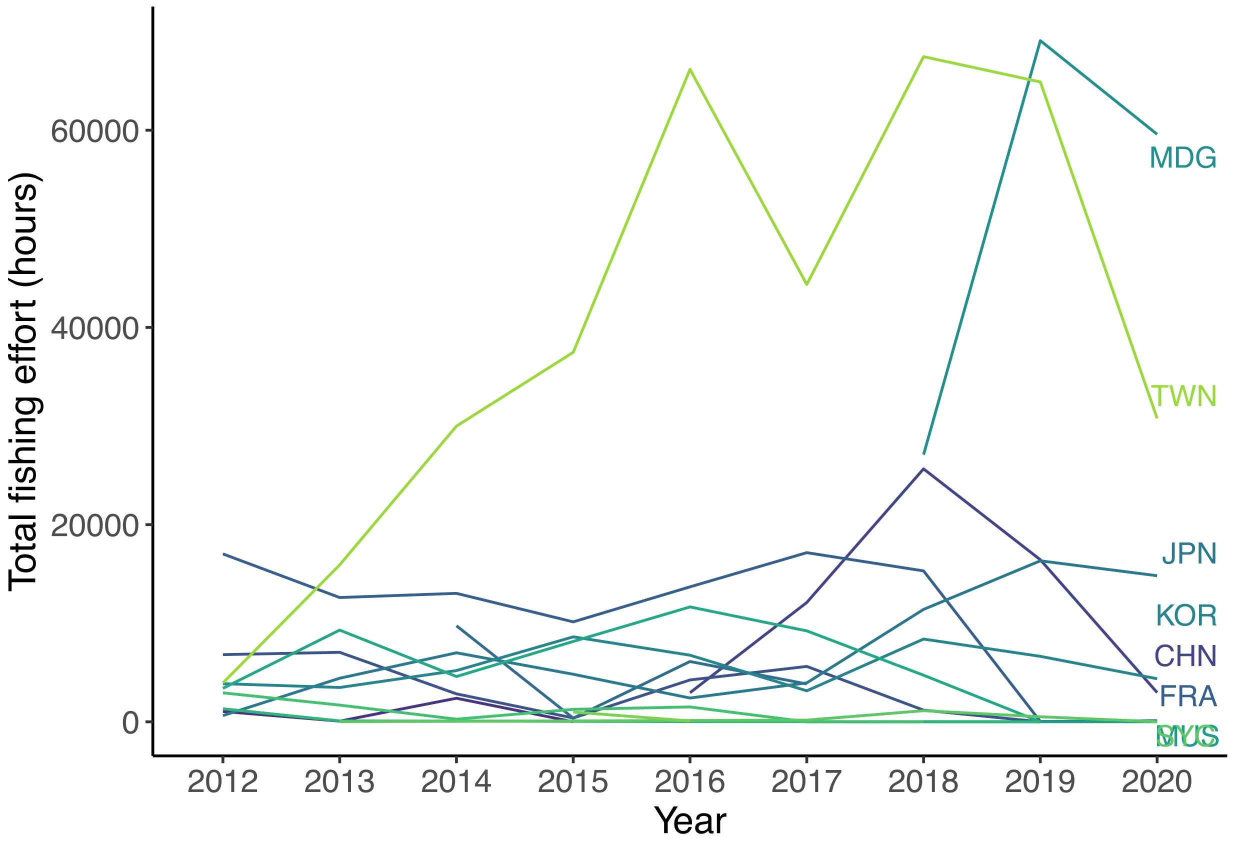


Figure S2: Total fishing effort (hours) for each country over time.

### Covariates

#### Oil Prices

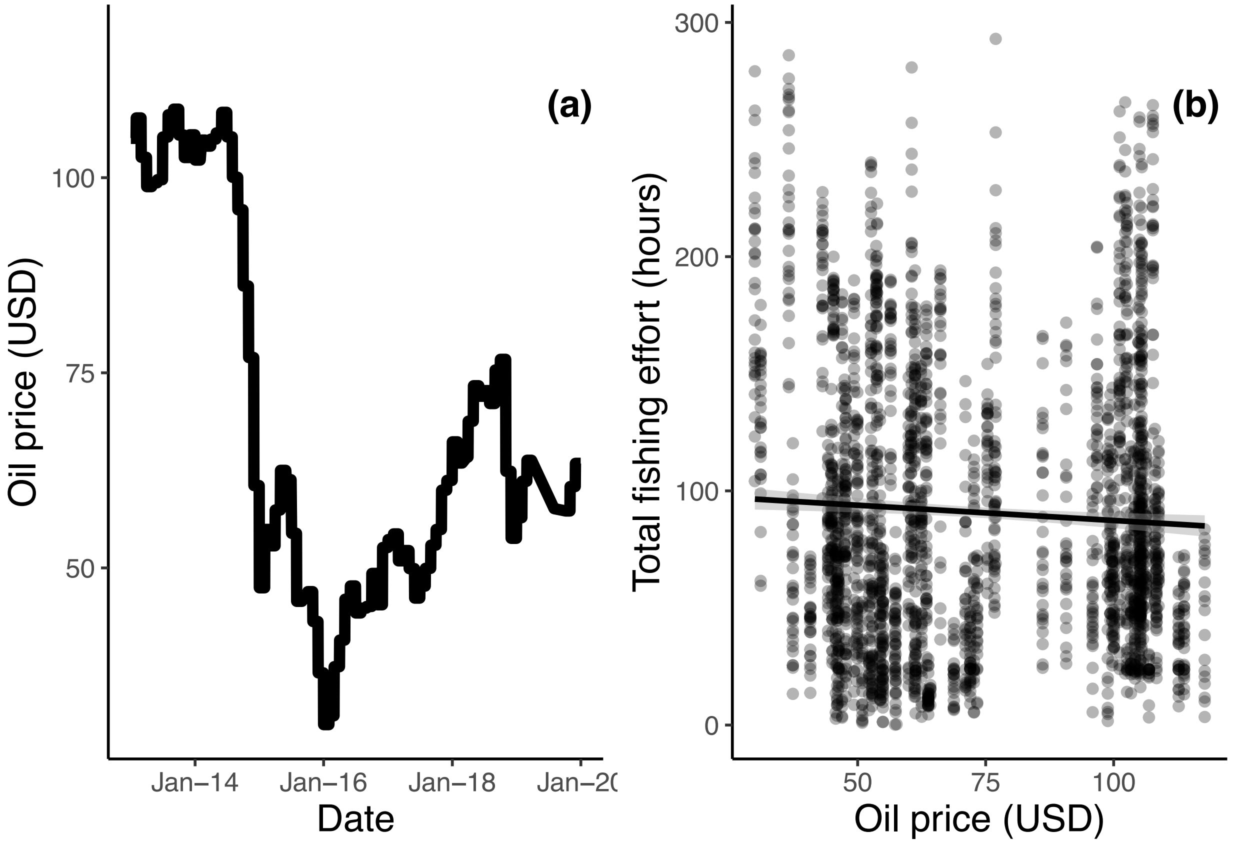


Figure S3: (a) Global oil price (USD) over time and (b) standarized fishing effort versus the global oil price.

#### Fish prices

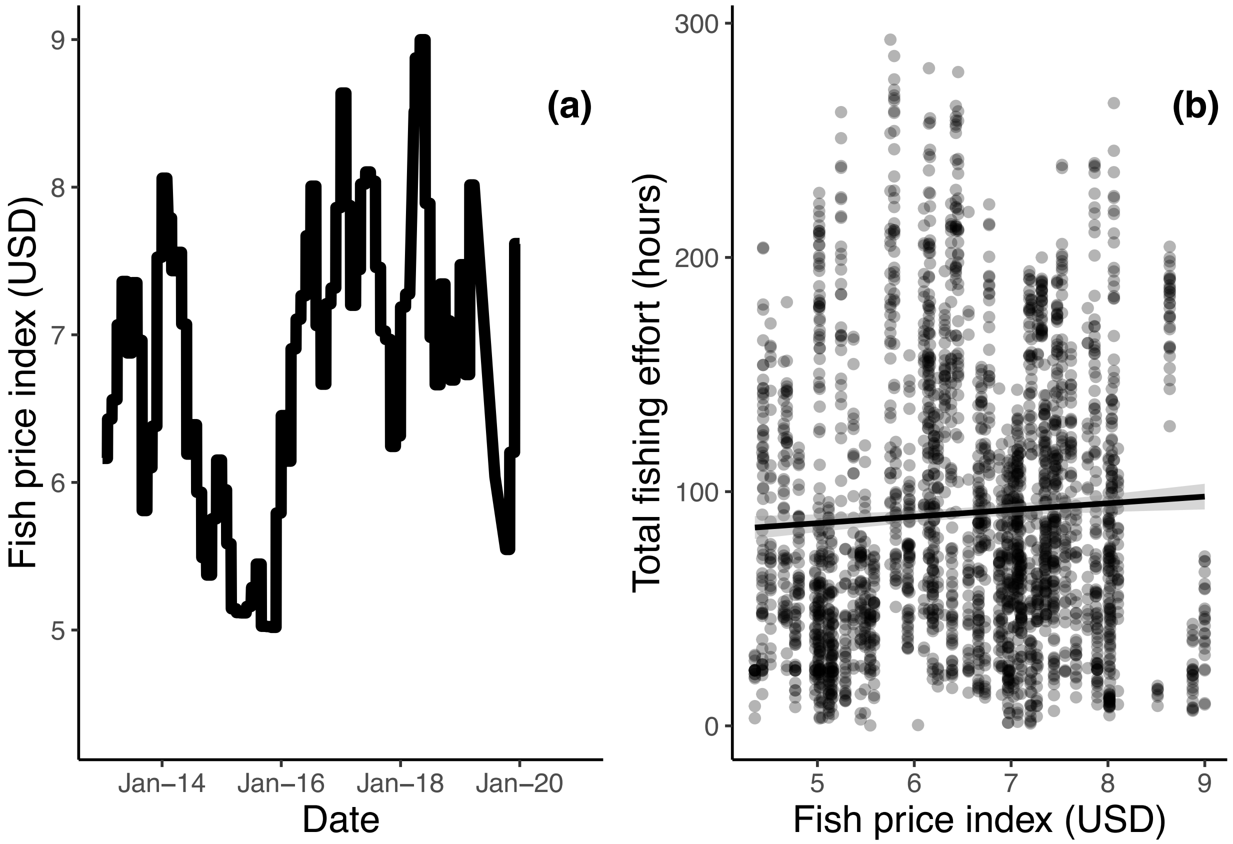


Figure S4: (a) Fish price index (USD) over time and (b) standarized fishing effort versus the fish price index.

#### Cyclone events

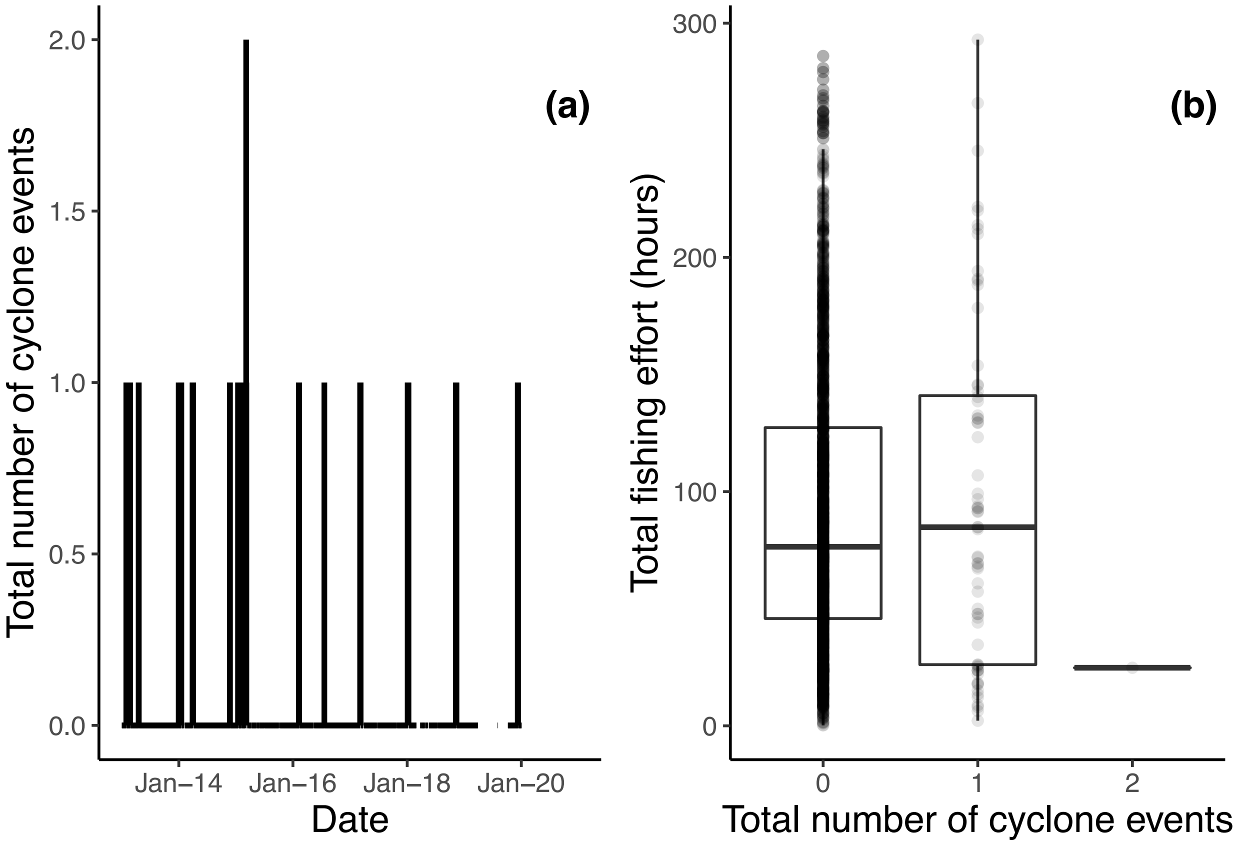


Figure S5: (a) Total number of cyclone events over time and (b) standarized fishing effort versus the total number of cyclone events.

#### Dipole index

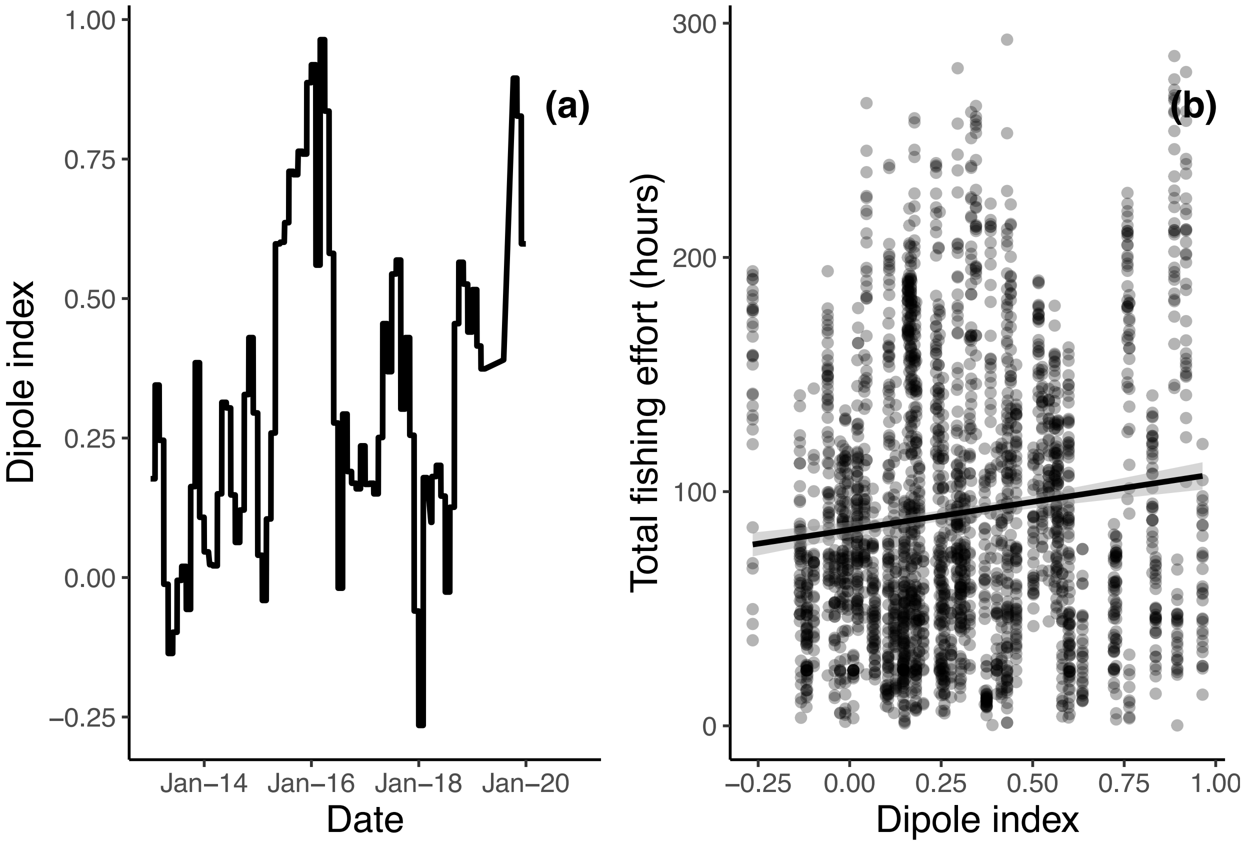


Figure S6: (a) Dipole index over time and (b) standarized fishing effort versus the dipole index.

### Diagnostic plots of best fitting model

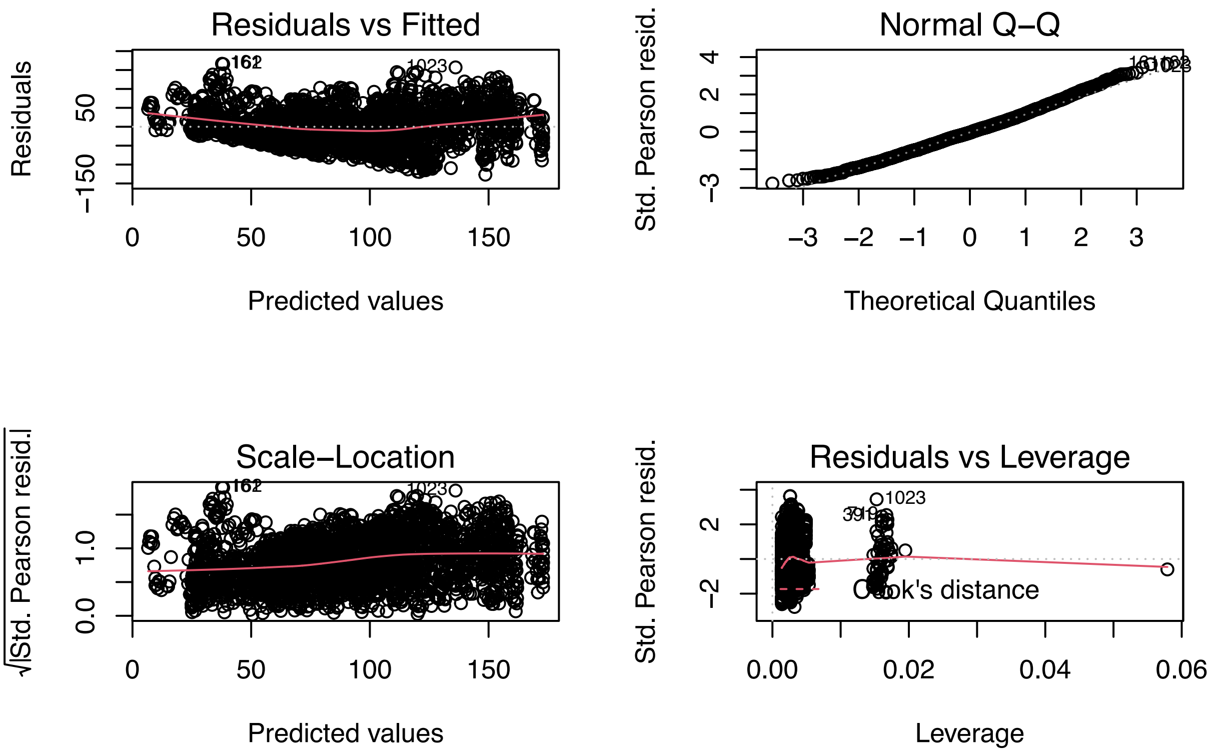


Figure S7: Diagnostic residual plots for the best fitting linear model described in the main text.

### Alternative models

In the main text, in our model selection process, we examined the total fishing effort for 2012-2019. Vessels were added to the Global Fishing Watch databases throughout the study making it difficult to evaluate which years of data to include or which vessels. Therefore, we present three additional analyses here with response variables as: fishing effort from 2017-2019, standardized fishing effort from 2017-2019, and only looking at data for vessels that were present every year.

Table S1: Parameter estimates (top value) and standard errors (bottom value in parentheses) for the average of the best fitting models (AIC < 2) using a GLM framework with a Gaussian error structure. Fishing effort (from 2017-2019) is the total daily fishing effort within the Madagascar EEZ divided by the cumulative number of vessels observed. Sine and cosine terms denote seasonal variables as a function of the julian day of the year.

| | Fishing effort |
|---|---|
| (Intercept) | $-28643.889$ |
| | (27779.515) |
| sine | $-119.333^{***}$ |
| | (14.267) |
| cosine | $326.734^{***}$ |
| | (8.804) |
| year | 14.229 |
| | (13.771) |
| Dipole index | $359.311^{***}$ |
| | (38.526) |
| Fish price index (USD) | $-33.968^{**}$ |
| | (12.018) |
| Oil price (USD) | $7.374^{***}$ |
| | (1.123) |
| Cyclone event | $-7.736$ |
| | (23.317) |
| Num. obs. | 1094 |

$^{***}p < 0.001$; $^{**}p < 0.01$; $^{*}p < 0.05$

Table S2: Parameter estimates (top value) and standard errors (bottom value in parentheses) for the average of the best fitting models (AIC < 2) using a GLM framework with a Gaussian error structure. Fishing effort (from 2017-2019) is the total daily fishing effort within the Madagascar EEZ divided by the cumulative number of vessels observed. Sine and cosine terms denote seasonal variables as a function of the julian day of the year.

| | Standardized fishing effort |
|---|---|
| (Intercept) | $301.582^{**}$ |
| | (116.688) |
| sine | $-0.696^{***}$ |
| | (0.057) |
| cosine | $1.686^{***}$ |
| | (0.046) |
| year | $-0.149^{**}$ |
| | (0.058) |
| Dipole index | $0.759^{***}$ |
| | (0.185) |
| Fish price index (USD) | $-0.007$ |
| | (0.034) |
| Oil price (USD) | $0.027^{***}$ |
| | (0.005) |
| Cyclone event | $-0.453^{*}$ |
| | (0.202) |
| Num. obs. | 1094 |

$^{***}p < 0.001$; $^{**}p < 0.01$; $^{*}p < 0.05$

Table S3: Parameter estimates (top value) and standard errors (bottom value in parentheses) for the average of the best fitting models (AIC < 2) using a GLM framework with a Gaussian error structure. Fishing effort (from 2012-2019) is the total daily fishing effort within the Madagascar EEZ for all vessels (as opposed to only the vessels seen during all nine years of the data set used in the main manuscript). Sine and cosine terms denote seasonal variables as a function of the julian day of the year.

| | Fishing effort |
|---|---|
| (Intercept) | $-121906.271^{***}$ |
| | (4774.274) |
| sine | $-95.190^{***}$ |
| | (4.954) |
| cosine | $213.735^{***}$ |
| | (4.662) |
| year | $60.584^{***}$ |
| | (2.373) |
| Dipole index | $110.033^{***}$ |
| | (14.962) |
| Fish price index (USD) | $-10.980^{**}$ |
| | (3.936) |
| Oil price (USD) | $1.556^{***}$ |
| | (0.199) |
| Cyclone event | $-50.379^{*}$ |
| | (19.549) |
| Log Likelihood | $-19061.089$ |
| AIC | 38140.179 |
| Delta | 0.000 |
| Weight | 1.000 |
| Num. obs. | 2897 |

$^{***}p < 0.001$; $^{**}p < 0.01$; $^{*}p < 0.05$

#### Madagascar MPA database

Table S4: Name and year started of areas in Madagascar listed as either a MPA or LMMA

| Name | Status.Year |
|---|---|
| Manombo | 1962 |
| Lokobe | 1966 |
| Nosy Tanihely | 1968 |
| Mananara Nord | 1990 |
| Baie de Baly | 1997 |
| Masoala Marine Park | 1997 |
| Tampolo Marine Park | 1997 |
| Tanjona Marine Park | 1997 |
| Nosy Ve | 1998 |
| Sahamalaza Iles Radama | 2001 |
| Loky Manambato | 2005 |
| Complexe Zones Humides Mahavavy Kinkony | 2006 |
| Velondriake | 2006 |
| Antongil Bay | 2007 |
| Nosy Hara | 2007 |
| Ranobe Bay | 2007 |
| Ambohibola | 2008 |
| Ambondrolava Mangroves | 2008 |
| Beheloke | 2008 |
| Itampolo | 2008 |
| Maromena/Befasy | 2008 |
| Tahosoa | 2008 |
| Amboditangena | 2009 |
| Ambodivahibe | 2009 |
| Antisakivolo | 2009 |
| Fimihara/Ankaranjelita | 2009 |
| Imorona | 2009 |
| Maintimbato | 2009 |
| Manjaboaka | 2009 |
| Rantohely | 2009 |
| Vohitralanana | 2009 |
| Ambodiforaha | 2011 |
| Analanjahana | 2011 |
| Aniribe | 2011 |
| Anoromby/Andreba | 2011 |

| | |
|---|---|
| Mahasoa | 2011 |
| Tanandava | 2011 |
| Teariake | 2011 |
| Tsinjoriake | 2011 |
| Vatolava | 2011 |
| Ambodimangamaro | 2012 |
| Anandrivola | 2012 |
| Hoalampano | 2012 |
| Seranambe | 2012 |
| Antanambe-Malotrandro | 2013 |
| Sainte Luce | 2014 |
| Ambato Atsinanana | 2015 |
| Ankarea | 2015 |
| Ankivonjy | 2015 |
| Elodrato | 2015 |
| Soariake | 2015 |
| Itapera | 2016 |
| Kirindy Mite | 2016 |
| Iles Barren | 2017 |
| Site Bioculturel dAntrema | 2017 |
| Nosy Ve Androka | 2018 |
